# How structural brain network topologies associate with cognitive abilities in a value-based decision-making task

**DOI:** 10.1101/743245

**Authors:** Cristina Bañuelos, Timothy Verstynen

**Affiliations:** Dept. Psychology, Carnegie Mellon University, Pittsburgh, Pennsylvania 15213; Dept. Biological Sciences, Carnegie Mellon University, Pittsburgh, Pennsylvania 15213; Carnegie Mellon Neuroscience Institute, Carnegie Mellon University, Pittsburgh, Pennsylvania 15213; Biomedical Engineering, Carnegie Mellon University, Pittsburgh, Pennsylvania 15213

**Keywords:** Value-based decision-making, Adaptive decision-making, Decision-making, Iowa Gambling Task, Graph-theoretic Topology, Structural brain networks

## Abstract

Value-based decision-making relies on effective communication across disparate brain networks. Given the scale of the networks involved in adaptive decision-making, variability in how they communicate should impact behavior; however, precisely how the topological pattern of structural connectivity of individual brain networks influences individual differences in value-based decision-making remains unclear. Using diffusion MRI, we measured structural connectivity networks in a sample of community dwelling adults (N=124). We used standard graph theoretic measures to characterize the topology of the networks in each individual and correlated individual differences in these topology measures with differences in the Iowa Gambling Task. A principal components regression approach revealed that individual differences in brain network topology associate with differences in optimal decision-making, as well as associate with differences in each participant’s sensitivity to high frequency rewards. These findings show that aspects of structural brain network organization can constrain how information is used in value-based decision-making.

**Abbreviations:** MRI - Magnetic Resonance Imaging; IGT – Iowa Gambling Task; DWI – Diffusion Weighted Imaging; QSDR – Q-Space Diffeomorphic Reconstruction; PCA – Principal Components Analysis; GLM – Generalized Linear Models

## Introduction

The brain is a highly organized network consisting of approximately 86 billion interconnected neurons (Herculano-Houzel, 2009). There are input-output computations made across all regions of the brain that are connected by bundles of axon fibers that communicate across long distances (Hopfield, 1982; Mountcastle, 1997). Fast and efficient communication throughout the brain is necessary for nearly all cognitive processes. This highly complex network is organized through cell bodies, dendrites, and axon terminals of these neurons that, together, make up the “grey matter,” whereas the axons connecting the cell bodies to the axon terminals make up the “white matter.” From a graph perspective, grey matter are the nodes that process information, and white matter forms the edges that determine which information is sent between nodes. The exact nature of the wiring architecture of the human brain, much like a circuit in a computer, impacts brain function, leading to subsequent cognition (McCulloch, 1944; Johansen-Berg, 2010; Hermundstad et al., 2014). The static organization of the structural architecture of the brain is both modular and hierarchical, supporting executing local operations and global integration of segregated functions (Park & Friston, 2013). Being able to measure the individual differences in structural connectivity of brain networks and their subsequent behavior would allow for the ability to explain the neural constraints on complex cognition (Verstynen, 2015).

The structural connectivity of brain networks is measured through a technique known as diffusion-weighted imaging (DWI). DWI takes advantage of the diffusion properties of water molecules within the axons of the myelinated white matter fascicles. Diffusion tensor imaging is one of the most popular DWI sampling schemes, sampling a few dozen orthogonal diffusion directions that are used to calculate a tensor of average diffusion direction within each voxel (for review, see Vettel et al., 2017). DWI has been used in conjunction with graph theoretic structural topology measures in order to understand the functional organization underlying structural networks (for review see Bullmore & Sporns, 2009).

There is growing evidence that brain networks have small-worldness properties, that can be characterized as a dense local clustering between neighboring nodes forming modules paired with short path length between any pairs between modules (Watts & Strogatz, 1998). This small-worldness supports the distributed nature of distinct brain areas while also demonstrating how these modules are integrated into global brain networks (Bassett & Bullmore 2006). However, we still have a limited understanding on how the topological organization of structural connections in the brain predicts individual differences in complex cognitive abilities.

In order to gain insights into how the brain may be organized to carry out executive processes, we looked into how the structural network organization may explain differences in executive abilities that are necessary to complete complex cognitive tasks. Using DWI methods, we measured whether individual differences in white matter topology, as seen through graph theoretic measures, associate with value-based decision-making (payoff or sensitivity to frequency of reward). Value-based decision-making is a complex task that, at a minimum, uses visual perception, attention, working memory, reinforcement learning, executive control, and other lower order functions in order to synthesize our decisions, and therefore relies on the efficient communication across global brain networks (Bechara et al., 1994). If small-worldness is a property of efficient network communication, then we hypothesize that individuals with more small world structural networks would be better at feedback driven, value-based decision-making.

## Material and Methods

### Participant Characteristics

We used an already collected sample of community dwelling adults in Pittsburgh, taken from the Weight-loss Intervention for brain Networks Project, collected at the University of Pittsburgh and Carnegie Mellon University. The sample of participants consisted of 124 participants (97 females, 27 males) between the ages of 22 and 55 (M= 44.38 years, SD=8.49). The participants had between 9 and 25 years of education (M= 16 years, SD=2.67). This research project was approved by the institutional review boards at both the University of Pittsburgh and Carnegie Mellon University.

### Iowa Gambling Task

Participants completed a computerized version of the IGT in which they are asked to select a card from any of the four decks presented with a varying amount of reward or punishment (Figure 1; Bechara et al., 1994). The participants were given a loan of $2000 and were instructed that the goal of the task is to maximize profit. They were also allowed to switch between any of the decks, at any time, and as often as they wished. The participants were not aware of any of the deck specifications and were only informed that each deck was different (Table 1). From each selection from Decks A or B (the “disadvantageous decks”), participants have a net loss of money. From each selection from Decks C or D (the “advantageous decks”), participants have a net gain of money. The amount of reward or punishment varied between decks and the position within a deck. Deck A and Deck B both had the same amount of overall net loss. However, in Deck A the punishment was more frequent and lower in magnitude, while in Deck B the punishment was less frequent, and higher in magnitude. Similarly, Deck C and Deck D had the same overall net gain. In Deck C the punishment was more frequent and smaller in magnitude, while Deck D is less frequent and higher in magnitude. From the selections made by the participants, their overall payoff (P=(C + D) − (A + B)) and their overall sensitivity to frequency of reward (Q= (B + D) − (A + C)) was calculated.

**Figure 1:**
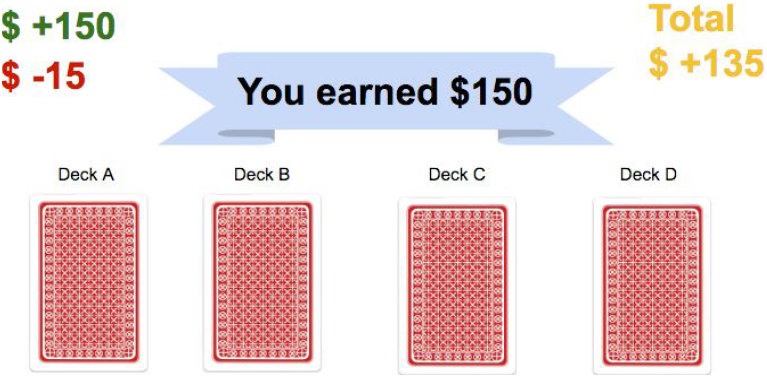
Computerized version of the Iowa Gambling Task. The computerized version of the IGT that participants were asked to complete is shown above. They were asked to select a card from any of the four decks presented with a varying amount of reward or punishment. The amount of cash earned and lost is shown in the green and red values, respectively.

**Table 1:** Iowa Gambling Task Deck Specifications. The decks have a different combination of two separate features including reward amount, frequency of reward, and result in either a net yield of overall loss or gain.

|  | A | B | C | D |
| --- | --- | --- | --- | --- |
| Reward | \$100 | \$100 | \$50 | \$50 |
| Frequency of Reward | Low | High | Low | High |
| Net Yield | Loss | Loss | Gain | Gain |

### Diffusion Weighted Imaging

All diffusion weighted images were acquired using a single-shell, diffusion tensor imaging protocol (voxel size = 2.4×2.4×2.4 mm, TR = 11100 ms, TE = 96 ms, 56 different directions, b-value = 2000 s/mm^2^). DWI data was reconstructed using a q-space diffeomorphic reconstruction (QSDR) method that creates models of water diffusion patterns for every voxel in the brain in an averaged space that allows for comparing across participants (Yeh & Tseng, 2011). A deterministic fiber tractography approach (Yeh et al., 2013) was then used to estimate the structural connections between brain areas for each participant (left in Figure 2), using standard tractography parameters (anisotropy threshold = 0.70, random seed orientation, tracked streamline count = 250,000, step size = 1mm, max turning angle = 75 degrees, smoothing = 0.50, minimum length = 20mm, maximum length = 160mm). The fiber tractography output was integrated with a gray matter brain atlas, using a well-established parcellation of distinct regions known as nodes (middle in Figure 2) to produce a connectivity matrix (right in Figure 2).

**Figure 2:**
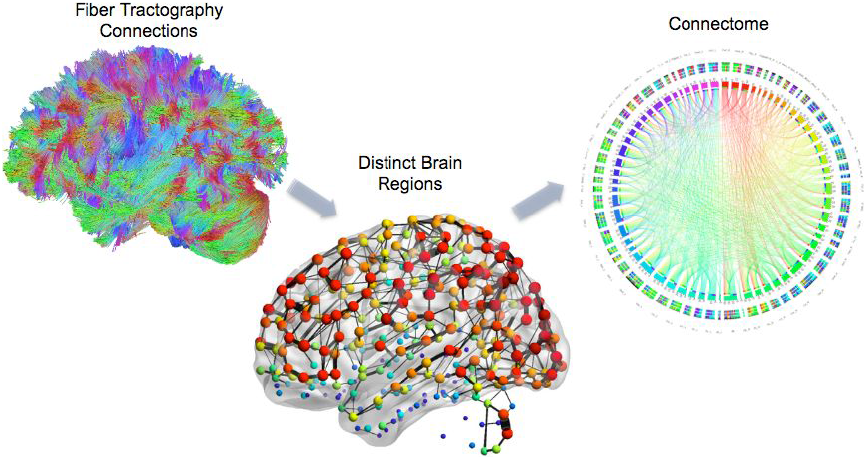
Brain Network Connectome. The fiber tractography DWI structural connections are used as the edges. They are then added to gray matter that is parcellated into distinct regions with a brain atlas to form the brain topologies. Together they form a brain graph representation of a brain network, and can be plotted as a connectome.

### Brain Network Connectivity Mapping

The weight of edges in the matrix were defined by the amount of streamlines connecting each pair of nodes (Tang et al., 2017). The analyses were also replicated using a different definition for edge weights in which they were equal to the number of individual white matter tracts connecting each pair of nodes divided by the total volume of the node pair. The network system was represented by a graph consisting of (*n*, *e*), where *n* refers to the node also known as the grey matter regions of interest, and *e* referred to the set of all edges on the graph. Each node was assigned with a real value. The node states present under one gray matter region were used to define the map. The structural topology measures (Figure 3) on each subject’s connectivity matrix include:

- <u>Density:</u> the fraction of present connections to all possible connections without taking into account any connection weights in the calculation (Rubinov & Sporns 2010)
- <u>Clustering Coefficient:</u> the fraction of triangles around a node (Rubinov & Sporns 2010)
- <u>Transitivity:</u> the ratio of triangles to triplets in the network, which can be used as an alternative measure to the clustering coefficient (Rubinov & Sporns 2010), although these are not identical metrics
- <u>Characteristic Path Length:</u> the average shortest path length in the network (Rubinov & Sporns)
- <u>Small-worldness:</u> dense local clustering or cliquishness of connections between neighboring nodes yet a short path length between any (distant) pair of nodes due to the existence of relatively few long-range connections (Bassett & Bullmore 2006)
- <u>Global Efficiency:</u> the average inverse shortest path length in the network (Rubinov & Sporns 2010)
- <u>Local Efficiency:</u> the global efficiency computed on node neighborhoods, and is related to the clustering coefficient (Rubinov & Sporns 2010)
- <u>Assortativity Coefficient:</u> a correlation coefficient between the degrees of all nodes on two opposite ends of a link, where a positive value would indicate that nodes tend to link to other nodes with the same or a similar degree (Rubinov & Sporns 2010)

**Figure 3:**
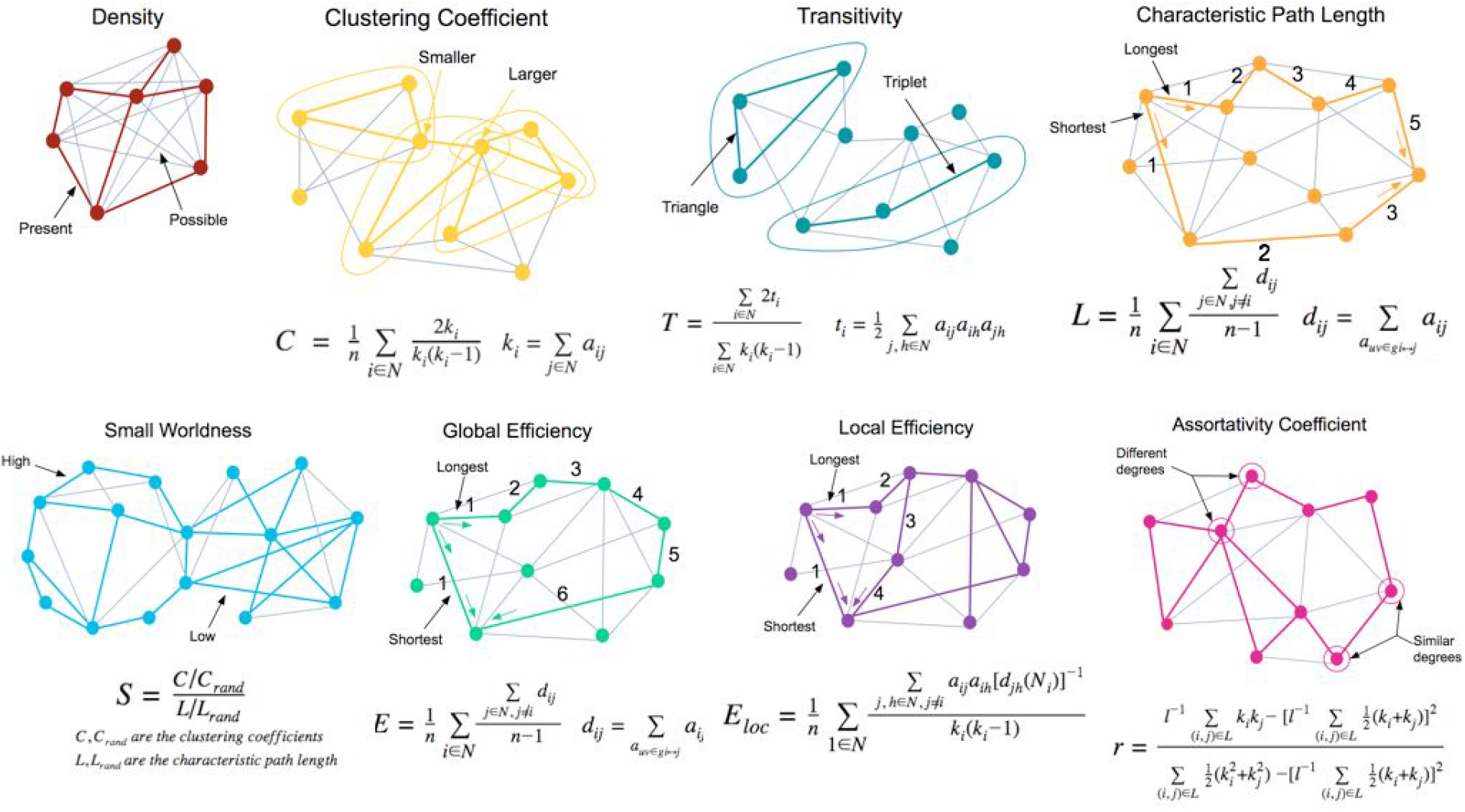
Graphed Structural Topology Measures. For each subject we looked at 8 measures of structural network topology, related to the “small worldness” of the brain networks (Rubinov & Sporns 2010; Bassett & Bullmore 2006). The measures included Density, Clustering Coefficient, Transitivity, Characteristic Path Length, Small Worldness, Global Efficiency, Local Efficiency, and Assortativity Coefficient.

## Results

### Characteristics and Distribution of Measures

The distribution measures calculated are minimum, quartile 1, median, quartile 3, maximum, skew, and the number of Grubbs test outliers (Table 2). The Grubbs test outliers were calculated with a significance alpha of 0.05. Transitivity, Characteristic Path Length, and Q scores have the greatest magnitude of skewness calculated with 1.074, 1.238, and −0.780, respectively. With the Grubbs test for calculating outliers, all of the variables had a total of 1 outlier.

**Table 2:** Distribution of Measures. The distribution measures calculated are minimum, quartile 1, median, quartile 3, maximum, skew, and the number of Grubbs test outliers.

|  | Minimum | Quartile.1 | Median | Mean | Quartile.3 | Maximum | Skew | Outliers |
| --- | --- | --- | --- | --- | --- | --- | --- | --- |
| Density | 0.042 | 0.052 | 0.055 | 0.055 | 0.058 | 0.07 | 0.027 | 1 |
| Clustering Coefficient | 0.128 | 0.226 | 0.253 | 0.253 | 0.277 | 0.389 | 0.113 | 1 |
| Transitivity | 0.005 | 0.023 | 0.036 | 0.039 | 0.053 | 0.128 | 1.074 | 1 |
| Net. Char. Path Length | 2.979 | 3.331 | 3.455 | 3.516 | 3.682 | 4.582 | 1.238 | 1 |
| Small Worldness | 0.036 | 0.063 | 0.072 | 0.072 | 0.081 | 0.106 | 0.011 | 1 |
| Global Efficiency | 0.296 | 0.337 | 0.349 | 0.348 | 0.36 | 0.396 | -0.359 | 1 |
| Local Efficiency | 9.367 | 16.878 | 18.656 | 18.67 | 20.631 | 28.049 | 0.078 | 1 |
| Assortativity | -0.308 | -0.162 | -0.085 | -0.079 | 0.008 | 0.191 | 0.15 | 1 |
| P Scores | -54 | 0 | 22 | 21.592 | 46 | 70 | -0.211 | 1 |
| Q Scores | -60 | 14 | 38 | 31.352 | 52 | 84 | -0.78 | 1 |

When looking across the network topology measures we found a high degree of correlation r across individuals (Figure 4). The greatest positive correlations observed were between Small Worldness and Local Efficiency with 0.88, Clustering Coefficient and Small Worldness with 0.90, and Clustering Coefficient and Local Efficiency with 0.97. Such strong correlations among the structural topology measures suggests that they share the common variance. High correlations among variables provides evidence of a low dimensional structure in the topology measures.

**Figure 4:**
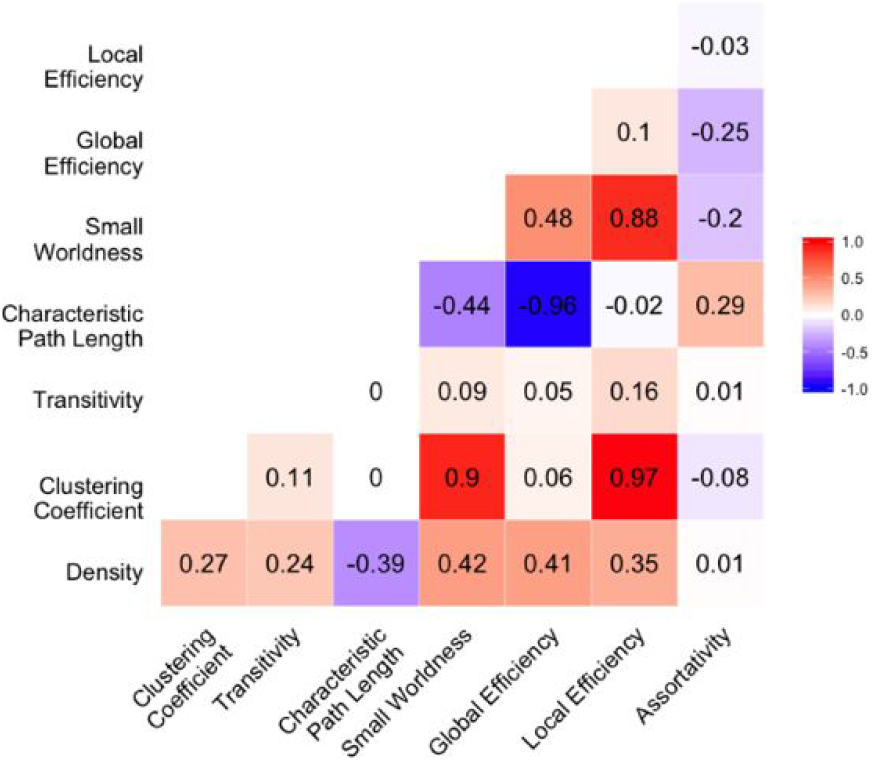
Correlation matrix of the structural topology measures. Strong negative correlations are depicted as blue, zero correlations are depicted as white, and strong positive correlations are depicted as red. Characteristic Path Length and Global Efficiency was found to have a strong negative correlation. Strong positive correlations were found between Small Worldness and Local Efficiency, Clustering Coefficient and Small Worldness, and Clustering Coefficient and Local Efficiency.

Since strong correlations were observed among the network topology measures in the data, we used Principal Component Analysis (PCA) to identify the lower dimensional components that explain most of the variance in these measures. PCA orthogonally transforms the highly correlated data into linearly uncorrelated principal components. PCA has the ability to reduce the dimensionality of data while maintaining useful information.

Table 4 shows the recovered 8 components and their mapping to the variables measured. Loadings were considered to be strong with a magnitude greater than 0.3. In the first PCA component all of the 8 variables measured are significantly loaded and there are strong positive loadings with Density, Clustering Coefficient, Small Worldness, and Local Efficiency, and a strong negative loading with Characteristic Path Length. The second component has significant loadings from 7 components, and has strong positive loadings with Global Efficiency, and strong negative loadings with Clustering Coefficient, Characteristic Path Length, and Local Efficiency. The third component has 5 variables with significant loadings and there is strong positive loadings with Density, Transitivity, and Assortativity, and has no strong negative loadings. The fourth component has 5 variables with significant loadings, and has a strong positive loading with Assortativity, and a strong negative loading with Transitivity. The fifth component has 6 variables with significant loadings, and it has a strong positive loading with Density, and strong negative loadings with Transitivity and Assortativity. The sixth component has 5 variables with significant loadings, with no strong positive loadings, and with strong negative loadings with Characteristic Path Length, Global Efficiency, and Local Efficiency. The seventh component has 5 variables with significant loadings, with strong positive loadings with Clustering Coefficient, Characteristic Path Length, Small Worldness, and Global Efficiency, and with a strong negative loading with Local Efficiency. The eighth component has 4 variables with significant loadings, with a strong positive loading with Clustering Coefficient, and a strong negative loading with Small Worldness. It is important to note that the last 3 components (6-8) have a similar pattern of loadings strength of loadings, with some variations in the sign of the loadings.

**Table 4:** Table of PCA Component Loadings for Variables. The second component has 7 measures with significant loadings, and it has a strong positive loading with Global Efficiency, and strong negative loadings with Clustering Coefficient, Characteristic Path Length, and Local Efficiency. The fifth component has 6 variables with significant loadings, and it has a strong positive loading with Density, and strong negative loadings with Transitivity and Assortativity.

| Components | 1 | 2 | 3 | 4 | 5 | 6 | 7 | 8 |
| --- | --- | --- | --- | --- | --- | --- | --- | --- |
| *Density* | 0.324 | 0.129 | 0.473 | 0.167 | 0.789 |  |  |  |
| *Clustering Coefficient* | 0.435 | -0.405 | -0.148 |  |  | 0.212 | 0.336 | 0.68 |
| *Transitivity* | 0.111 |  | 0.702 | -0.618 | -0.327 |  |  |  |
| *Characteristic Path Length* | -0.302 | -0.565 |  | -0.111 | 0.209 | -0.527 | 0.469 | -0.187 |
| *Small Worldness* | 0.525 | -0.114 | -0.146 |  | -0.135 | 0.285 | 0.322 | -0.696 |
| *Global Efficiency* | 0.33 | 0.529 |  | 0.122 | -0.269 | -0.633 | 0.333 | 0.115 |
| *Local Efficiency* | 0.447 | -0.389 |  |  |  | -0.436 | -0.67 |  |
| *Assortativity* | -0.132 | -0.229 | 0.487 | 0.748 | -0.361 |  |  |  |

When looking at the percent variance accounted for (Figure 5), we found that only the first 5 components explained 95% of the data. Therefore our subsequent analysis only focuses on these components.

**Figure 5:**
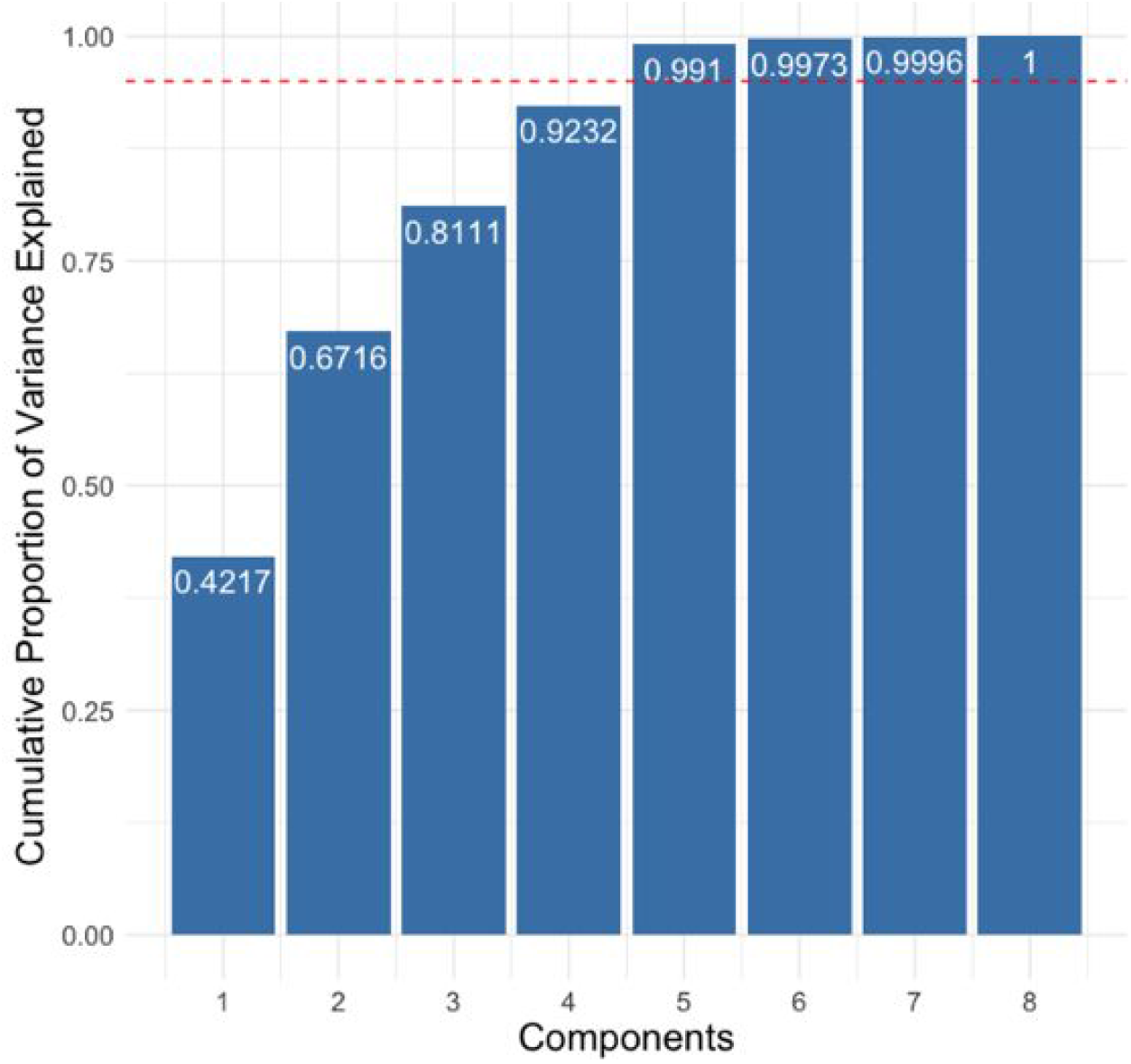
Principal Components for 95% Variance Explained. The cumulative proportion variance for each of the 8 PCA components are plotted in the blue bars. The red dotted line demonstrates the mark for 95% of cumulative proportion of variance. The barplot shows that components 1 through 5 account for over 95% of the cumulative proportion variance. Components 1-5 have a cumulative proportion variance that is about equal to 99.10%.

### Associations with IGT Performance

We used the generalized linear model (GLM) to identify which of the 5 principal components that explained the most variance were associated with the P and Q scores of the participants. We were able to find a significant association between the second component and the P score, which reflects the ability to effectively use feedback to maximize returns, with a significance level of less than 0.1 (Table 5). The GLM with the P scores and the first 5 principal components predicts P scores with a correlation of 0.232. The rest of the components do not appear to be able to predict P scores accurately with a significance level of less than 0.1. The second component was found to have mean bootstrap coefficient estimate of 3.05 and a lower and upper bound of 2.735 and 3.365 (Table 9).

**Table 5:** GLM Analysis on Principal Components to Predict P Score. The generalized linear model to predict P Scores with the PCA components shows that the second component was able to reliably predict P scores with a significance of p < 0.1.

|  | Coefficient Estimate (Std. Error) |
| --- | --- |
| Intercept | 21.83(2.51)*** |
| Component 1 | 1.72(1.37) |
| Component 2 | 3.07(1.77) . |
| Component 3 | 0.99(2.38) |
| Component 4 | -2.36(2.66) |
| Component 5 | 3.07(3.42) |

Significant codes:

Furthermore, components 1, 2, 3, and 5 were found to have positive coefficient estimates with standard errors ranging from 1.37 to 3.42 (Table 5). Component 4 was found to have negative coefficient estimate with standard error of 1.77 (Table 5). Component 2 with a coefficient estimate of 3.07 and standard error of 1.77 was the only component found to can predict P Scores with a significance of 0.1 (Table 5). The Intercept was found to have a coefficient estimate of 21.83 and a standard error of 2.51 with a significance of p < 0.001 (Table 5). This is lower than the Bonferroni corrected threshold of 0.005 (corrected for 10 comparisons). The cross validation errors calculated from the GLM are 1.06 and 1.06, which are significantly low values. The rest of the components do not appear to be able to predict P scores accurately with a significance level of less than 0.1.

In addition, we saw a significant association between the fifth component and the Q score, which reflects a sensitivity to high frequency rewards (Table 6) with a significance of p < 0.05. The GLM with the Q scores and the first 5 principal components predicts Q scores with a correlation of 0.318. The rest of the components do not appear to be able to predict Q scores accurately with a significance level of less than 0.1. The fifth component was found to have mean bootstrap coefficient estimate of 7.579 and a lower and upper bound of 6.7 and 8.065 (Table 9).

**Table 6:** GLM Analysis on Principal Components to Predict Q Score. Component 5 with a coefficient estimate of 7.46 and standard error of 3.52 was the only component found to can predict Q Scores with a significance of 0.01. The rest of the components do not appear to be able to predict Q scores with a significance level of less than 0.01.

|  | Coefficient Estimate (Std. Error) |
| --- | --- |
| Intercept | 31.33(2.59)*** |
| Component 1 | -0.77(1.41) |
| Component 2 | 2.25(1.83) |
| Component 3 | -3.48(2.46) |
| Component 4 | 2.91(2.74) |
| Component 5 | 7.58(3.52)* |

Furthermore, components 2, 4, and 5 were found to have positive coefficient estimates with standard errors ranging from 1.41 to 3.52 (Table 6). Components 1 and 3 were found to have negative coefficient estimates with standard errors of 1.41 and 2.46, respectively (Table 6). Component 5 with a coefficient estimate of 7.46 and standard error of 3.52 was the only component found to can predict Q Scores with a significance of 0.01 (Table 6). The Intercept was found to have a coefficient estimate of 31.44 and a standard error of 2.59 with a significance of p < 0.001 (Table 6). This is lower than the Bonferroni corrected threshold of 0.005 (corrected for 10 comparisons). The cross validation errors calculated from the GLM are 1.06 and 1.06, which are significantly low values. The rest of the components do not appear to be able to predict Q scores accurately with a significance level of less than 0.1.

Using only the second component model coefficients as weights for each component in predicting P Scores, we can transform the value of the weights from the components to data space with the topology measures. Their data space transformation demonstrates that Global Efficiency leads to an increase in P Score, whereas Characteristic Path Length, Clustering Coefficient, Global Efficiency, Local Efficiency, and Assortativity lead to a decrease in P Score. Global Efficiency has the greatest positive effect on P score, and Characteristic Path Length has the greatest negative effect (Table 7).

**Table 7:** Structural Topology Weights of Component 2 in Topology Space. In topology space, component 2 was made up of primarily Characteristic Path Length, Global Efficiency, Local Efficiency, Transitivity, Clustering Coefficient, and Assortativity.

|  | Weight | SE of Weight | Lower Bound | Upper Bound |
| --- | --- | --- | --- | --- |
| *Density* | 0.396 | 0.23 | -0.055 | 0.847 |
| *Clustering Coefficient* | -1.242 | 0.721 | -2.656 | 0.172 |
| *Transitivity* | -0.192 | 0.112 | -0.411 | 0.027 |
| *Characteristic Path Length* | -1.732 | 1.006 | -3.703 | 0.239 |
| *Small Worldness* | -0.348 | 0.202 | -0.744 | 0.048 |
| *Global Efficiency* | 1.62 | 0.941 | -0.224 | 3.464 |
| *Local Efficiency* | -1.192 | 0.692 | -2.549 | 0.165 |
| *Assortativity* | -0.701 | 0.407 | -1.5 | 0.097 |

Using only the fifth component model coefficients as weights for each component in predicting Q Scores, we can transform the value of the weights from the components to data space with the topology measures. Their data space transformation demonstrates that Density and Characteristic Path Length lead to an increase in Q Score, whereas Clustering Coefficient, Transitivity, Small Worldness, Global Efficiency, Local Efficiency, and Assortativity lead to a decrease in Q Score (Table 8). Density has the greatest positive effect on Q score, and transitivity has the greatest negative effect.

**Table 8:** Structural Topology Weights of Component 5 in Topology Space. In topology space, component 5 was made up of primarily Density, Transitivity, Global Efficiency, & Assortativity.

|  | Weight | SE of Weight | Lower Bound | Upper Bound |
| --- | --- | --- | --- | --- |
| *Density* | 5.9827 | 2.7808 | 0.5323 | 11.4331 |
| *Clustering Coefficient* | -0.4699 | 0.2184 | -0.8981 | -0.0418 |
| *Transitivity* | -2.476 | 1.1509 | -4.7316 | -0.2203 |
| *Characteristic Path Length* | 1.5864 | 0.7374 | 0.1411 | 3.0317 |
| *Small Worldness* | -1.0268 | 0.4773 | -1.9623 | -0.0914 |
| *Global Efficiency* | -2.0363 | 0.9465 | -3.8914 | -0.1812 |
| *Local Efficiency* | -0.2837 | 0.1319 | -0.5422 | -0.0252 |
| *Assortativity* | -2.738 | 1.2726 | -5.2323 | -0.2436 |

**Table 9:** Bootstrapping Analysis on Principal Components with P and Q Scores. Component 2 with P Score was found to have a mean bootstrap coefficient estimate of 3.05, whereas Component 5 with Q Score was found to have a mean bootstrap coefficient estimate of 7.579.

|  | Component Estimate | Mean Bootstrap | Standard Deviation | Error | Lower Bound | Upper Bound |
| --- | --- | --- | --- | --- | --- | --- |
| *Component 2 with P Score* | 3.065 | 3.05 | 1.791 | -0.315 | 2.735 | 3.365 |
| *Component 5 with Q Score* | 7.579 | 7.382 | 3.877 | -0.682 | 6.7 | 8.065 |

## Discussion

Here we tested the hypothesis that individuals with more small-world structural brain networks would be better at feedback driven value-based decision-making. First, we found that the graph topology measures of white matter networks had a low dimensional structure that could be mostly explained by five principal components. A regression analysis examining how these components correlated with the ability to use feedback to maximize long term payoffs (P) and the sensitivity to high frequency rewards (Q) found associations with both. One component, component 2, mapping positively on Global Efficiency, and negatively on Characteristic Path Length and Local Efficiency topology measures, reliably associated with the ability to use feedback to maximize rewards, such that the positive weights associate with greater ability, and the negative weights associate with lesser ability (Table 7). Another component, component 5, mapping positively on Density and Characteristic Path Length, and negatively on Transitivity, Global Efficiency, and Assortativity topology measures, reliably associated with sensitivity to rewards, such that these positive weights associate with greater sensitivity, and negative weights associate with a lesser sensitivity (Table 8).

Since the strength of connections in white matter networks is highly plastic (Yeh et al., 2016), a more optimal design may have been a longitudinal study in which individuals were tracked over time and compared to themselves. Furthermore, we did not look at node-wise topology measures, and primarily focused on using the topology measures of global networks. It could be that small differences in specific pathways, such as the cortico-basal ganglia thalamic loops (Alexander, DeLong, & Strick, 1986), are highly predictive of aspects of value-based decision-making, but by looking at global networks this effect is washed out. Future directions could include focusing on regions of interest, or subnetworks of interest, in order to identify individual network associations rather than global network associations. In addition, the functional dynamics of the global networks could be more sensitive to individual differences in decision-making. By looking at functional connectivity or task-linked responses, instead of structural connectivity, we may be able to understand the nature of the communication across these networks in order to understand their associations.

Future applications of this work include using the topological organization of the structural connectivity of brain networks as neuroimaging markers for different decision-types or other cognitive states. For example, performing a similar analysis on other complex decision tasks (e.g., the Stroop task). In addition, future studies could be more strategic in the brain networks of interest. For example, future work should have a more detailed focus on regions of interest such as the basal ganglia pathways, that plays a critical role reinforcement learning and decision-making or integrate both structural and functional connectivity measures, to provide a holistic understanding of the network form and function.

In summary, we have shown here how the topology of white matter networks have a low dimensional structure in their topological organization and a subset of these low dimensional components reliably associated with individual differences in the ability to use feedback to modify future decisions.

## Acknowledgements

This research was supported through the mentorship of Timothy Verstynen, Ph.D., I would also like to appreciate the guidance from Alexis Porter, and Daniel Brasier, Ph.D in the completion of this research project.

